# Wear and Tear of the Intestinal Visceral Musculature by Intrinsic and Extrinsic Factors

**DOI:** 10.1101/2021.08.26.457849

**Authors:** Ho D. Kim, Eric So, Jiae Lee, Yi Wang, Vikram S. Gill, Anna Gorbacheva, Hee Jin Han, Katelyn G.-L. Ng, Ken Ning, Inez K.A. Pranoto, Alejandra J.H. Cabrera, Dae Seok Eom, Young V. Kwon

## Abstract

The gut visceral musculature plays essential roles in not only moving substances through the lumen but also maintaining the function and physiology of the gut. Although the development of the visceral musculature has been studied in multiple model organisms, how it degenerates is poorly understood. Here, we employ the *Drosophila* midgut as a model to demonstrate that the visceral musculature is disrupted by intrinsic and extrinsic factors, such as aging, feeding, chemical-induced tissue damage, and oncogenic transformation in the epithelium. Notably, we define four prominent visceral musculature disruption phenotypes, which we refer as ‘sprout’, ‘discontinuity’, ‘furcation’, and ‘crossover’ of the longitudinal muscle. Given that the occurrence of these phenotypes is increased during aging and under various stresses, we propose that these phenotypes can be used as quantitative readouts of deterioration of the visceral musculature. Intriguingly, administration of a tissue-damaging chemical dextran sulfate sodium (DSS) induced similar visceral musculature disruption phenotypes in zebrafish larvae, indicating that ingestion of a tissue-damaging chemical can disrupt the visceral musculature in a vertebrate as well. Our study provides insights into the deterioration of the gut visceral musculature and lays a groundwork for investigating the underlying mechanisms in *Drosophila* as well as other animals.

## Introduction

Developmental processes shape sophisticated adult tissue structures and patterns. These structures and patterns are maintained over time and also deteriorated by intrinsic and extrinsic factors, which may affect tissue physiology and organismal health. In *Drosophila*, the adult visceral musculature of the midgut comprises longitudinal and circular muscle fibrils, which form a well-defined lattice pattern (Amcheslavsky et al., 2009; Klapper, 2000). The final form of visceral muscle is shaped during metamorphosis and persists to sustain the function and physiology of the midguts in adults (Amcheslavsky et al., 2009; Klapper, 2000). During metamorphosis, the larval midgut undergoes extensive remodeling. Immediately after pupation, the larval visceral muscle loses its syncytial state and de-differentiates into mononucleated cells known as ‘secondary myoblasts’ (Amcheslavsky et al., 2009; Klapper, 2000). As metamorphosis of the midgut progresses, secondary myoblasts fuse back and differentiate to generate the adult syncytial musculature (Amcheslavsky et al., 2009; Klapper, 2000).

Progressive degeneration of tissue structures is a part of development. Intrinsic and extrinsic factors, such as extensive usage and diseases, can disrupt a tissue’s structure and pattern. Although the gut visceral musculature is likely to be degenerated during aging or in a certain disease context, it is poorly understood how the process happens in *Drosophila* as well as other animals. Degeneration of the visceral musculature can not only affect its physiology and function, but that of the gut and whole organism as well, an effect that is exemplified in a previous study that characterized the role of the *Drosophila* ortholog of *Mitochondrial Distribution and Morphology Regulator 1 misato* (*mst*) in the visceral muscle (Min et al., 2017). Depletion of *mst* disrupted the actin-myosin structure in the visceral muscle, which presumably impairs the visceral muscle function. Interestingly, depletion of *mst* in the muscle caused various phenotypes, including abnormal enlargement of midguts, reduction in gut mobility, defects in feeding, and decreased lifespan. Additionally, previous studies have shown that *Drosophila* midgut visceral muscle communicates with the midgut epithelium by secreting various signaling ligands, such as the *Drosophila* Wnt Wingless (Wg), *Drosophila* insulin-like peptide 3 (Dilp3), transforming growth factor-β Decapentaplegic (Dpp), and a neuregulin-like EGFR ligand Vein (Vn) (Biteau and Jasper, 2011; Guo et al., 2013; Jiang and Edgar, 2009; O’Brien et al., 2011; Scopelliti et al., 2014; Xu et al., 2011; Zhou et al., 2015). These myokines play essential roles in the regulation of intestinal stem cells (ISCs) during normal homeostasis and tissue damage. Therefore, degeneration of the visceral musculature during aging and under stress is likely to affect the function and physiology of the midgut epithelium.

In this study, we demonstrate that the midgut visceral musculature deteriorates during aging and under various stresses in *Drosophila*. In particular, we define four quantifiable visceral musculature disruption phenotypes and show that the occurrence of the phenotypes are affected by intrinsic and extrinsic factors, such as feeding, administration of tissue-damaging chemical, aging, and oncogenic transformation. Notably, administration of dextran sulfate sodium (DSS) induced similar visceral musculature disruption phenotypes in zebrafish, suggesting that chemical-induced tissue damage can also disrupt the visceral musculature of a vertebrate. Thus, our study not only provides insights into the deterioration of the visceral musculature and tools to study the underlying mechanisms in *Drosophila* but other organisms as well.

## Results

### Expression of mutant *Ras* (*Ras^V12^*) in intestinal epithelial cells disrupts the visceral musculature

Manipulation of the signaling pathways controlling ISC division can induce hyperplasia in the midgut epithelium . It has been shown that expression of a mutant *Ras* (*Ras^V12^*) in ISCs and enteroblasts (EBs) with *esg-GAL4*, *UAS-GFP*, *tub-GAL80^ts^* (referred as *esg^ts^*; see methods) induces midgut epithelial hyperplasia (Jiang and Edgar, 2009; Lee et al., 2020). However, these *Ras^V12^* cells quickly disappear from the midgut by apically delaminating into the lumen and basally disseminating into hemocoel (Lee et al., 2020). Notably, basally disseminating *Ras^V12^* cells produce actin- and cortactin-rich invasive protrusions, which are associated with degradation of the extracellular matrix (ECM) and the visceral muscle layer (Lee et al., 2020). Thus, expression of *Ras^V12^* with *esg^ts^* in adult midguts leads to a severe disruption of the visceral musculature, which is manifested by frequent discontinuation of longitudinal muscles in the posterior midguts (Figure 1A and 1B) (Lee et al., 2020). Given these observations, we leveraged the *Ras^V12^*-induced intestinal hyperplasia model to uncover phenotypes associated with the disruption of the visceral musculature.

**Figure 1:**
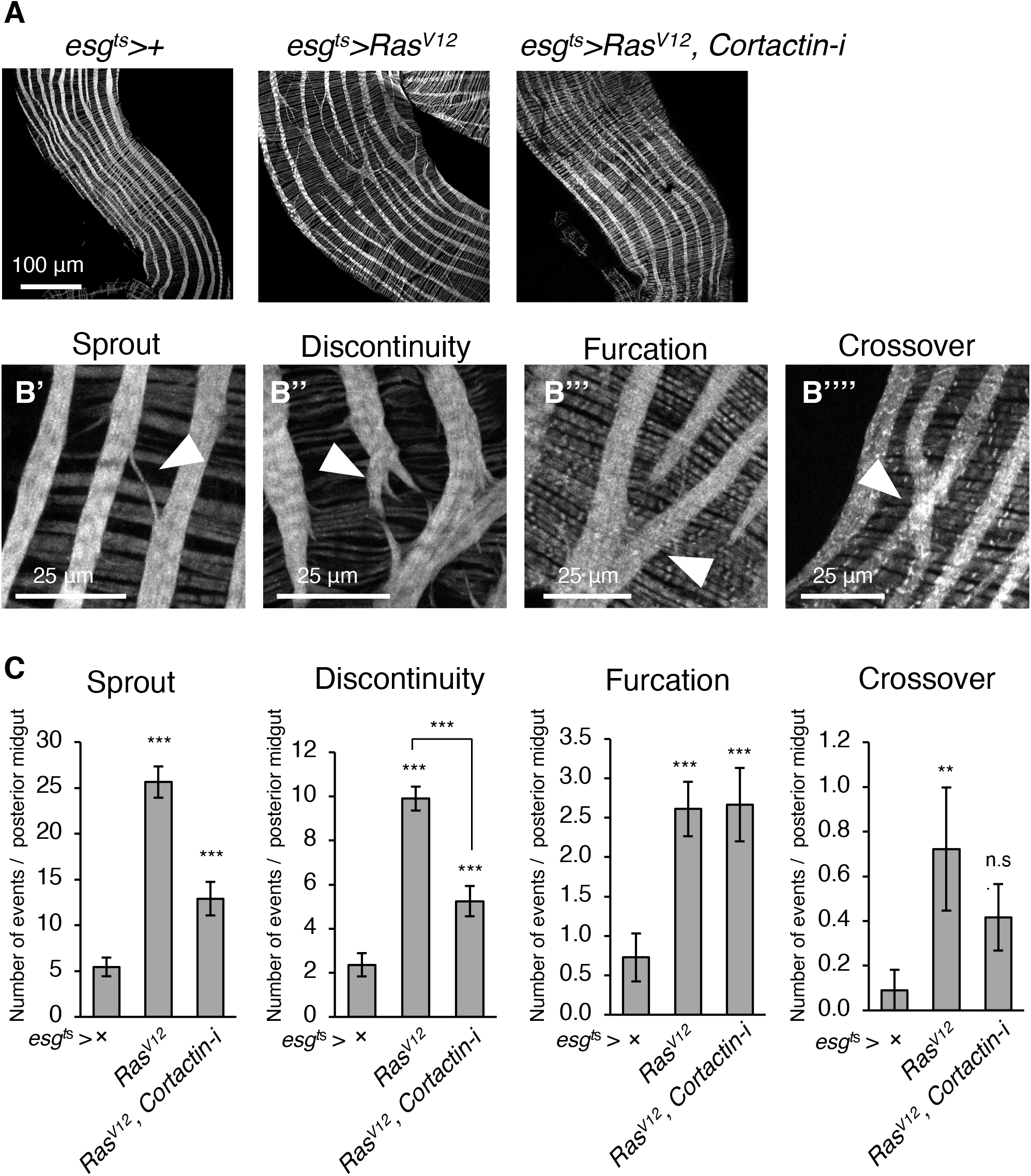
*Ras^V12^* expression in ISCs and EBs results in significant deterioration of the visceral muscle pattern. **A.** Representative images of the visceral musculature of the posterior midgut. Transgenes were induced for 2 days by shifting to 29°C. *HMS00658* was used to knockdown *Cortactin* (*Cortactin-i*). Visceral muscle was stained by phalloidin (gray). **B.** Four visceral musculature disruption phenotypes observed in the posterior midguts. The representative images of the phenotypes referred as ‘Sprout’ (**B’**), ‘Discontinuity’ (**B’’**), ‘Furcation’ (**B’’’**), and ‘furcation’ (**B’’’’**) are shown. **C.** Quantification of the visceral musculature disruption phenotypes. The data represent mean±SEMs of the indicated phenotype in the posterior midguts. N = 20 (*esg^ts^*); N = 20 (*esg^ts^*>*Ras^V12^*); N = 12 (esg^ts^>*Ras^V12^*, *Cortactin RNAi*) replicates. *p<0.05; **p<0.01; ***p<0.001 compared to *esg^ts^*>+ (control) unless indicated by bracket (Student’s t-test).

By analyzing the visceral muscle images of *esg^ts^>+* (control) and *esg^ts^>Ras^V12^* posterior midguts, we identified four prominent visceral musculature disruption phenotypes (Figure 1B). The first phenotype is ‘sprout’, which we define as a thin branch of muscle fiber emerging from the lateral side of longitudinal muscles (Figure 1B’). Sprouts are also observed in control midguts but they are more frequently detected in *esg^ts^*>*Ras^V12^* posterior midguts (Figure 1C). The second phenotype, which is referred as ‘discontinuity’, describes a pause of longitudinal muscle. Although most of the longitudinal muscles continuously span the posterior midgut without disruption (Figure 1A), on average about two longitudinal muscles are discontinuous in control midguts (Figure 1B’’ and 1C). These discontinuous longitudinal muscles are normally found at the anterior region of posterior midguts (Figure S1, arrowheads), suggesting that they could be generated as a part of normal developmental process. However, longitudinal muscle discontinuity is increased approximately five times in *esg^ts^*>*Ras^V12^* midguts compared to controls (Figure 1C). Additionally, the occurrence of discontinuity is not limited to the anterior region of *esg^ts^*>*Ras^V12^* posterior midguts (Figure S1). A significant portion of discontinuities in *esg^ts^*>*Ras^V12^* midguts are observed pairs (Figure S1), suggesting that they might be formed by longitudinal muscle breakage. The third phenotype is ‘furcation’ of longitudinal muscle, which is defined as one longitudinal muscle strand dividing into two or more smaller strands (Figure 1B’’’). When control and *esg^ts^*>*Ras^V12^* midguts are compared, furcation is detected 5 times more in *esg^ts^*>*Ras^V12^* midguts (Figure 1C). The fourth phenotype, which we refer as ‘crossover’ of longitudinal muscle, is characterized as X-shaped longitudinal muscle, which is formed when a longitudinal muscle strand goes over an adjacent longitudinal muscle strand (Figure 1B’’’’). Longitudinal muscle crossover is rarely detected in control midguts while the occurrence is significantly increased in *esg^ts^*>*Ras^V12^* posterior midguts (Figure 1C). Altogether, we have identified four visceral musculature disruption phenotypes by analyzing the visceral musculature in *esg^ts^*>*Ras^V12^* posterior midguts. These phenotypes are easily detectable and quantifiable.

Cortactin plays a key role in the function of invasive protrusions, which are formed at the basal side of *Ras^V12^*-expressing intestinal epithelial cells (Lee et al., 2020). To address whether the increased disruption of the visceral musculature in *esg^ts^*>*Ras^V12^* midguts was associated with the action of invasive protrusions, we depleted *Cortactin* in *Ras^V12^*-expressing intestinal epithelial cells to inhibit the function of invasive protrusions. Intriguingly, depletion of *Cortactin* significantly reduced the occurrence of sprout and discontinuity while the occurrence of furcation was not affected (Figure 1C). Note that we couldn’t discern whether crossover was a phenotype associated with the function of invasive protrusions in *esg^ts^*>*Ras^V12^* midguts because of their low occurrence rate. These observations suggest that alteration of neighboring intestinal epithelial tissue’s physiology can disrupt the visceral musculature. In particular, we show that a cellular process involved in cell invasion is partially responsible for the occurrence of sprout and discontinuity, not furcation in *esg^ts^*>*Ras^V12^* midguts. Additionally, these observations also suggest that these visceral musculature disruption phenotypes are not entirely interdependent.

### Intestinal epithelial hyperplasia induced by activation of Yorkie disrupts the visceral musculature

Hippo signaling keeps ISCs in check during normal homeostasis (Karpowicz et al., 2010; Ren et al., 2010; Shaw et al., 2010). Thus, inhibition of Hippo signaling or activation of the downstream transcription factor Yorkie leads to proliferation of ISCs, resulting in formation of *Drosophila* midgut epithelial hyperplasia (Kwon et al., 2015; Kwon et al., 2013; Kwon et al., 2019). To address how midgut expansion due to hyperplasia affects the visceral musculature, we expressed an active form of *yorkie* (*yki^3S/A^*) with *esg^ts^*, which induced progressive midgut epithelial hyperplasia (Figure 2A, 2B, and S2). We started to see a significant accumulation of *yki^3S/A^* cells at day 4, which became more prominent at later days (Figure S2). The occurrence of sprout was generally higher from the beginning of *yki^3S/A^* expression, suggesting that a slight change in the physiology of ISCs and EBs due to *yki^3S/A^* expression might be sufficient to increase sprouts. Interestingly, the sprout phenotype was more frequently detected at day 6 of *yki^3S/A^* expression. In contrast, the occurrence of discontinuity and furcation started to significantly increase at either day 4 or 6 of *yki^3S/A^* expression when *yki^3S/A^*-induced hyperplasia became prominent (Figure 2C and S2). Expression of *yki^3S/A^* with *esg^ts^* didn’t increase the occurrence of crossover above the background levels (Figure 2C).

**Figure 2:**
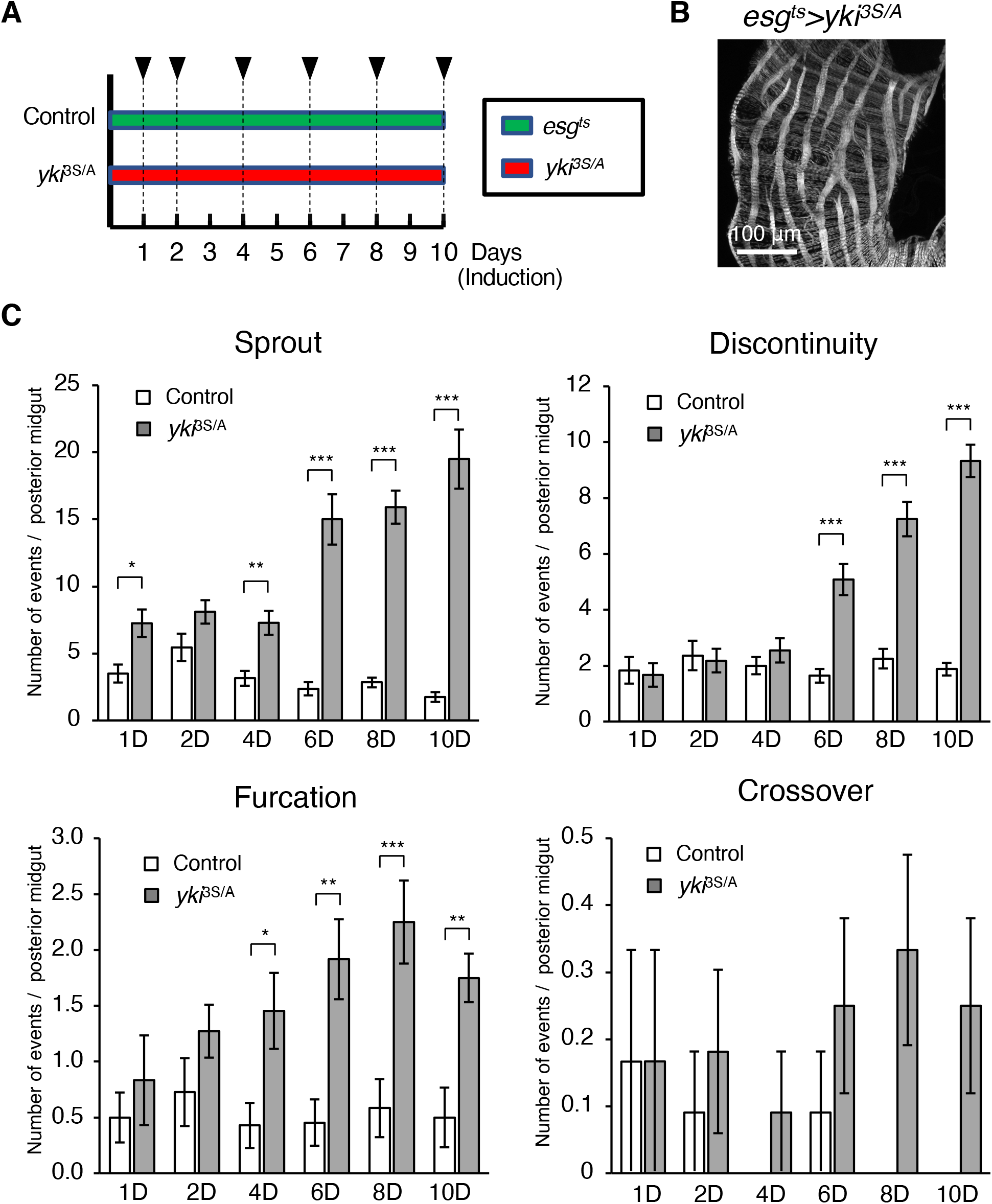
*yki^3S/A^* expression in ISCs and EBs disrupts the visceral musculature. **A.** The experimental schematic. Transgenes were induced by incubating flies at 29°C. Induction periods (days) are shown at the bottom. Arrowhead indicates time of dissection. **B.** Representative image of posterior midgut after 8 day of *yki^3S/A^* expression. Visceral muscle is stained by phalloidin (gray). **C.** Quantification of the visceral musculature disruption phenotypes. The represented data are mean±SEMs quantified in the posterior midguts. N = 6 (1 day, *esg^ts^*>+); 6 (1 day, *esg^ts^*>*yki^3S/A^*); 11 (2 day, *esg^ts^*>+); 11 (2 day, *esg^ts^*>*yki^3S/A^*); 7 (4 day, *esg^ts^*>+); 11 (4 day, *esg^ts^*>*yki^3S/A^*); 12 (6 day, *esg^ts^*>+); 12 (6 day, *esg^ts^*>*yki^3S/A^*); 12 (8 day, *esg^ts^*>+); 12 (8 day, *esg^ts^*>*yki^3S/A^*); 8 (10 day, *esg^ts^*>+); 12 (10 day, *esg*^ts^>*yki*^3S/A^) midguts. *p<0.05; **p<0.01; ***p<0.001 compared to the same day *esg^ts^*>+ (Student’s t-test).

These results indicate that midgut epithelial hyperplasia induced by expression of *yki^3S/A^* is sufficient to induce the visceral musculature disruption phenotypes, including sprout, discontinuity, and furcation. Given the correlation between the occurrence of these phenotypes and the magnitude of hyperplasia, we speculate that expansion of the midgut might be a factor causing the visceral musculature disruption phenotypes. Nevertheless, it might not be possible to attribute the cause of the visceral musculature disruption phenotypes to one factor since the occurrence of sprout is also increased even before a hyperplasia is formed in *esg^ts^*>*yki^3S/A^* midguts. Taken these results and the observations with *Ras^V12^* expression with *esg^ts^* together, we conclude that stresses associated with oncogenic transformation in the neighboring epithelial tissue can disrupt the visceral musculature.

### Feeding rich diet after fasting disrupts the visceral musculature

The four phenotypes resulting from disruption of the visceral musculature by oncogenic stress in the midgut epithelia can be used as quantitative readouts for assessing the visceral musculature disruption by intrinsic and extrinsic stressors. Given the speculation that expansion of the midgut epithelium can be a factor affecting the integrity of the visceral musculature, we designed an experiment to test the effects of starvation and re-feeding, which significantly changes the midgut size (O’Brien et al., 2011).

Midgut epithelium is a highly plastic tissue, which can quickly resize by adapting to the feeding state (O’Brien et al., 2011). Previous studies have shown that midguts significantly shrink under fasting and expand during feeding (O’Brien et al., 2011). ISCs and EBs in the midgut epithelium can contribute to tissue growth in adult flies. However, the visceral muscle doesn’t have stem or progenitor cells. Although sudden expansion or shrinkage of the midgut epithelium can lead to an alteration in overall tension in the visceral muscle, it is not understood whether changes in midgut size due to feeding or fasting can affect the integrity of the visceral musculature. To address this, we put flies under three different feeding conditions: 1) 4 days on a continuous rich diet, 2) 2 days on rich diet and then 2 days on water, and 3) 2 days on rich diet, 2 days on water, and then 2 days on rich diet (Figure 3A). Consistent with the previous observations, 2 days of fasting significantly reduced the posterior midgut width and re-feeding for 2 days after fasting allowed the midguts to grow back (Figure 3B). Interestingly, fasting for 2 days had no effect on the occurrence of the visceral musculature disruption phenotypes (Figure 3C). In contrast, re-feeding a rich diet after fasting significantly increased the occurrence of sprout and discontinuity when compared to either 2 days of feeding followed by 2 days fasting or 4 days of continuous feeding (Figure 3C). The occurrence of furcation and crossover was unaltered by any of the feeding conditions. These results suggest an interesting association between the midgut size plasticity and the occurrence of the visceral musculature disruption phenotypes. Midgut shrinkage (Figure 3B) due to fasting is not a factor causing disruption of the visceral musculature. Our observations suggest that sudden midgut expansion due to re-feeding after fasting can be a factor associated with disruption of the visceral musculature.

**Figure 3:**
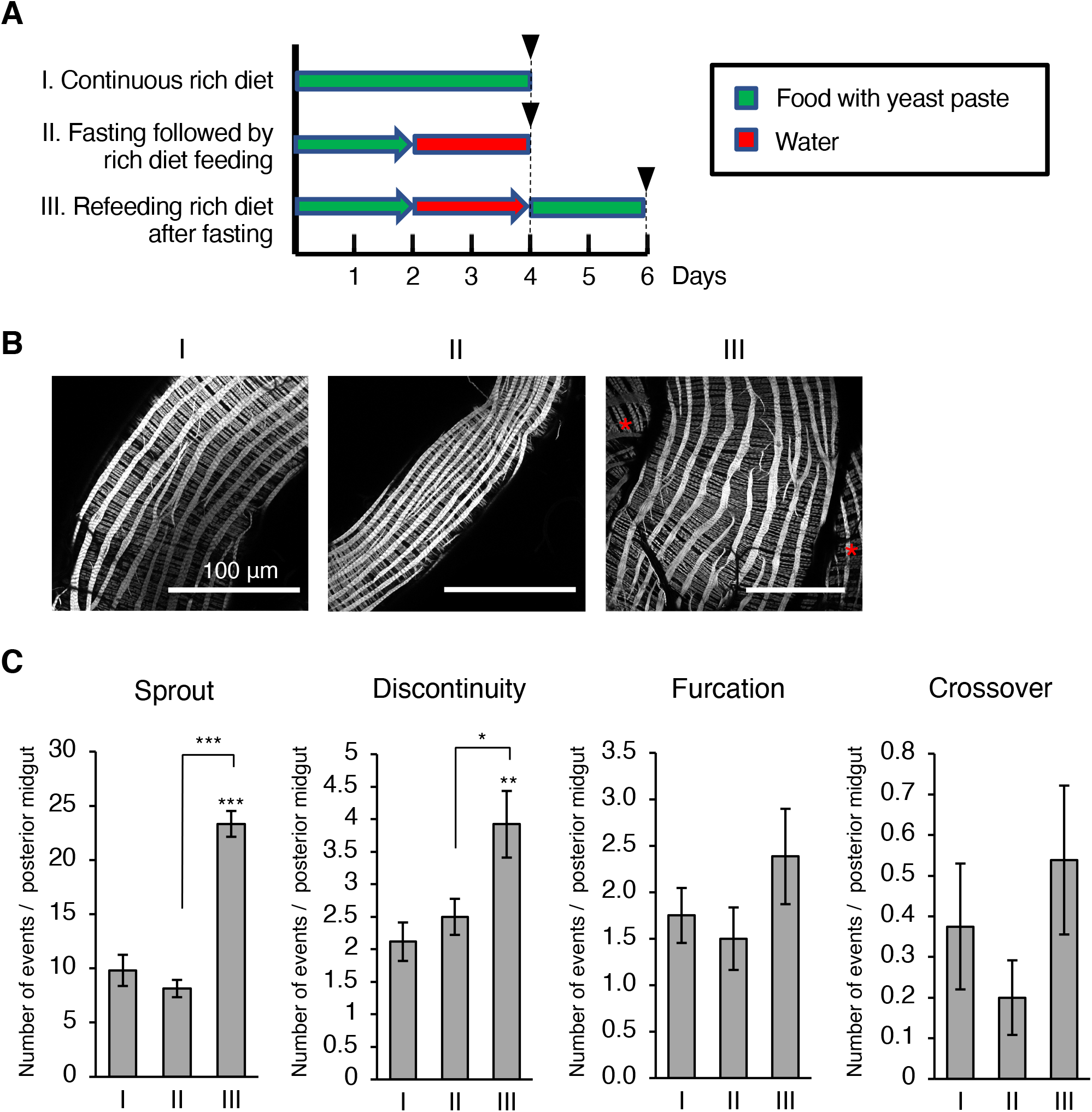
Feeding rich diet after fasting disrupts the visceral musculature. **A.** The experimental schematic. Arrowhead indicates time of dissection. **B.** The visceral musculatures of the posterior midgut. I) continuous rich diet (cornmeal food supplemented with yeast paste in 5% sucrose), II) rich diet and then fasting (water only), and III) re-feeding with rich diet after fasting. **C.** Quantification of the visceral musculature disruption phenotypes. Mean±SEMs are shown. N = 17 (Rich Diet only); 22 (Fasting); 13 (Fasting then re-feeding). *p<0.05; **p<0.01; ***p<0.001 compared to the corresponding condition I measurement unless indicated by bracket (Student’s t-test).

### Feeding bleomycin disrupts the visceral musculature

Ingestion of tissue-damaging agents or pathogenic bacteria induces injury to the midgut epithelium, which is followed by a tissue regeneration response (Amcheslavsky et al., 2009). During tissue damage and regeneration, the midgut epithelium significantly remodels as damaged cells are eliminated and ISCs proliferate to replace the damaged cells (Amcheslavsky et al., 2009; Ren et al., 2010; Staley and Irvine, 2010). In *Drosophila*, bleomycin has been commonly used to induce a tissue damage and regeneration response in midguts. To address the impact of chemical-induced tissue damage on the occurrence of the visceral musculature disruption phenotypes, we assessed how feeding bleomycin for 4 days in 5% sucrose solution altered the occurrence of the visceral musculature disruption phenotypes (Figure 4A). We found that bleomycin feeding significantly increased the occurrence of all the four visceral musculature disruption phenotypes (Figure 4B) compared to controls (5% sucrose only). Thus, we concluded that ingestion of tissue-damaging agents could be a factor that affected the integrity of the visceral musculature. Although the bleomycin-induced midgut epithelial damage and regeneration response has been well studied, it is not known if feeding bleomycin directly damages the visceral musculature. Thus, we cannot discern whether the visceral musculature disruption is caused indirectly by an interaction with the damaged midgut epithelium or directly by damage to the visceral musculature by bleomycin.

**Figure 4:**
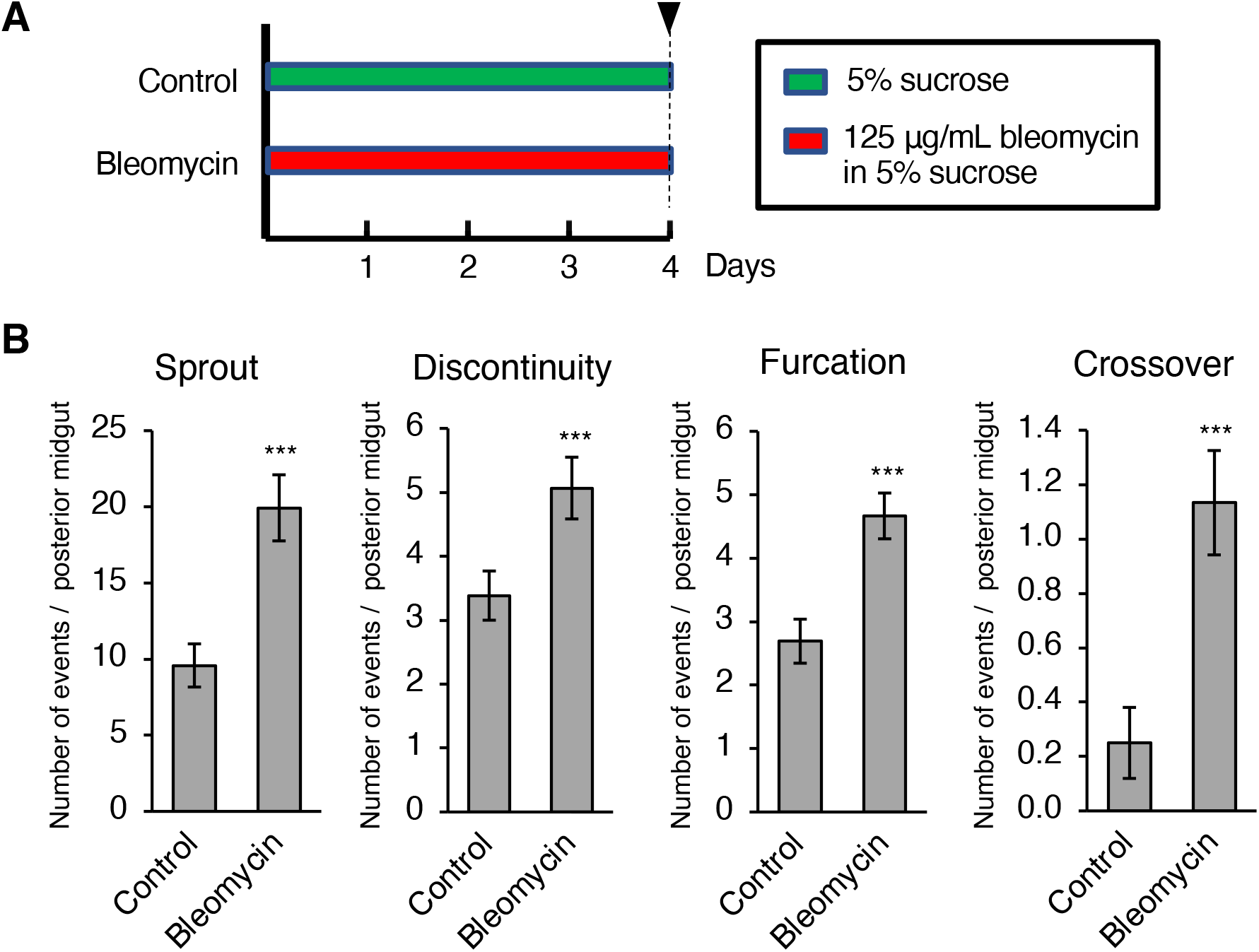
Bleomycin-induced tissue damage increases the visceral musculature disruption phenotypes. **A.** The experimental schematic. Arrowhead indicates time of dissection. **B.** Quantification of the visceral musculature disruption phenotypes. Mean±SEMs are shown. N = 13 (Sucrose); 15 (Bleomycin). *p<0.05; **p<0.01; ***p<0.001 compared to the corresponding 5% sucrose measurement (Student’s t-test).

### Aging is a factor associated with disruption of the visceral musculature

Deterioration of tissue integrity and function is a hallmark of aging (Fleckenstein et al., 2021; Gustafsson and Ulfhake, 2021; Jin, 2010; Li et al., 2013). Our findings demonstrate that various stressors can disrupt the visceral musculature. Thus, we hypothesized that aging could also be a factor causing the visceral musculature disruption phenotypes. To address how aging alters the visceral musculature, we compared the occurrence of the visceral musculature disruption phenotypes in 5-, 20-, and 60-day old flies (Figure 5A). Interestingly, we found that the occurrence of sprout progressively increased as flies age (Figure 5B). Additionally, a significant increase in the occurrence of discontinuity was observed in 60-day old flies (Figure 5B). The occurrence of furcation and crossover remained unchanged in 20- or 60-day old flies when compared to 5-day old flies (Figure 5B). These results indicate that the visceral musculature deteriorates during aging, which is manifested by the increased occurrence of sprout and discontinuity in old animals. The absence of changes in the furcation and crossover phenotypes suggests that aging might be the mildest stressor compared to other treatments in our study.

**Figure 5:**
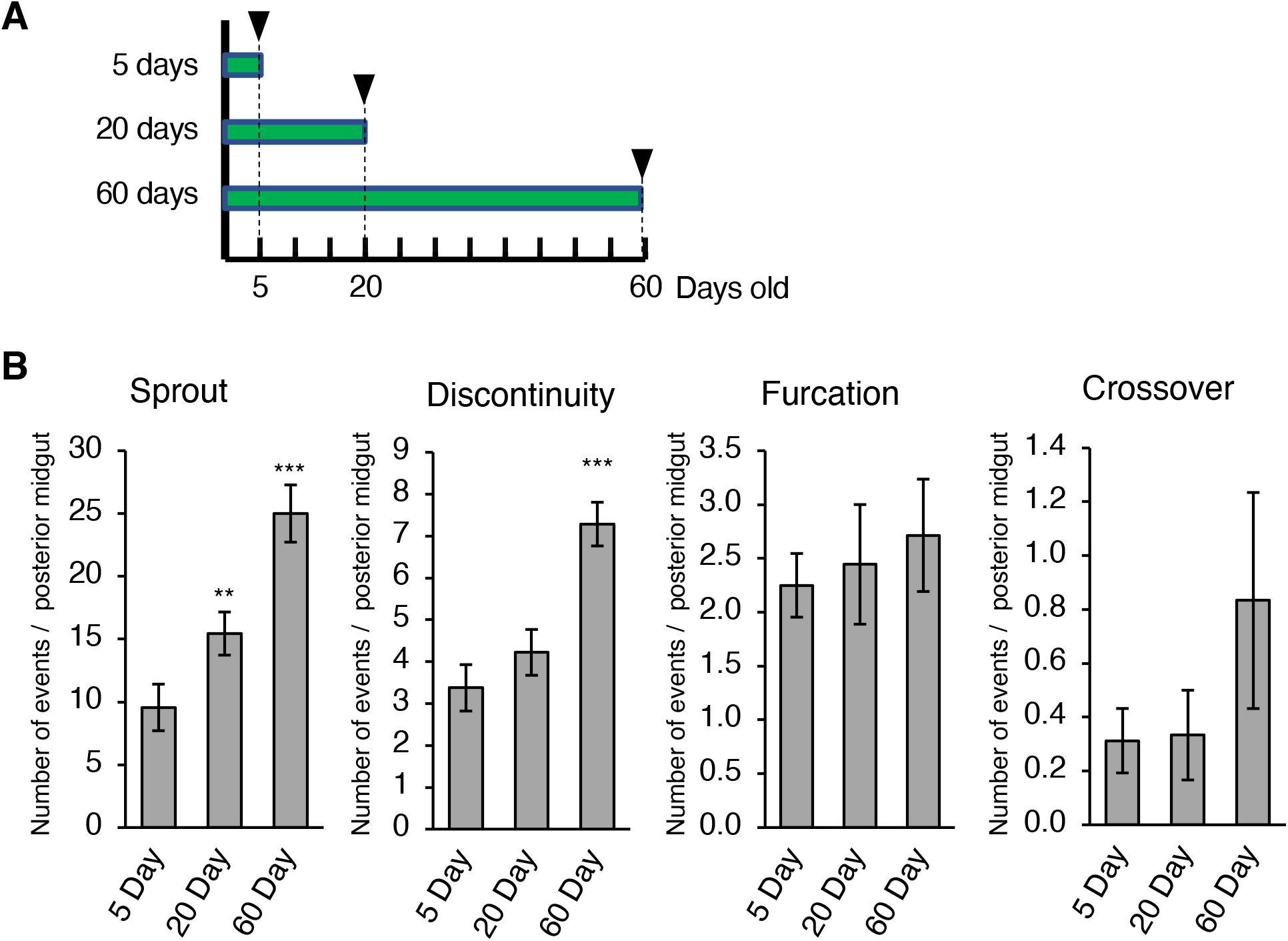
Aging is a factor associated with disruption of the visceral musculature. **A.** The experimental schematic. Arrowhead indicates time of dissection. **B.** Quantification of visceral musculature damage phenotypes per category. The data represents the mean ± S.E.M of specific phenotype per gut observed. N = 16 (5 day), N = 17 (10 day), and N = 7 (60 day). *p<0.05; **p<0.01; ***p<0.001 compared to the corresponding 5 day measurement (Student’s t-test).

### Similar visceral musculature disruption phenotypes are observed in zebrafish upon DSS administration

Our study revealed that the integrity of the visceral musculature can be compromised by various stressors in *Drosophila*. Given the lack of studies in other animals, we sought to address whether we can observe similar visceral musculature disruption phenotypes in *Danio rerio*. Dextran sulfate sodium (DSS) is a chemical commonly used for generation of chemically-induced intestinal damage in animal models (Chuang et al., 2019; Kiesler et al., 2015; Oehlers et al., 2013; Oehlers et al., 2017). In zebrafish larvae (Figure 6A), it has been shown that administration of DSS can recapitulate a few pathological features of human inflammatory bowel disease (IBD) (Chuang et al., 2019; Oehlers et al., 2013; Oehlers et al., 2017). Strikingly, administration of 1% DSS for 1 day affected the integrity of the visceral musculature in zebrafish larvae (Figure 6B and 6C). Aggregate-like phalloidin signals were frequently observed in the visceral musculature from DSS-treated animals while those signals were rarely detected in the visceral musculature from control animals (Figure 6D). In addition, we could spot the longitudinal muscle phenotypes resembling furcation and crossover specifically in the intestines from DSS-treated animals (Figure 6D). These observations demonstrate that chemical-induced disruption of the visceral musculature can also occur in a vertebrate.

**Figure 6:**
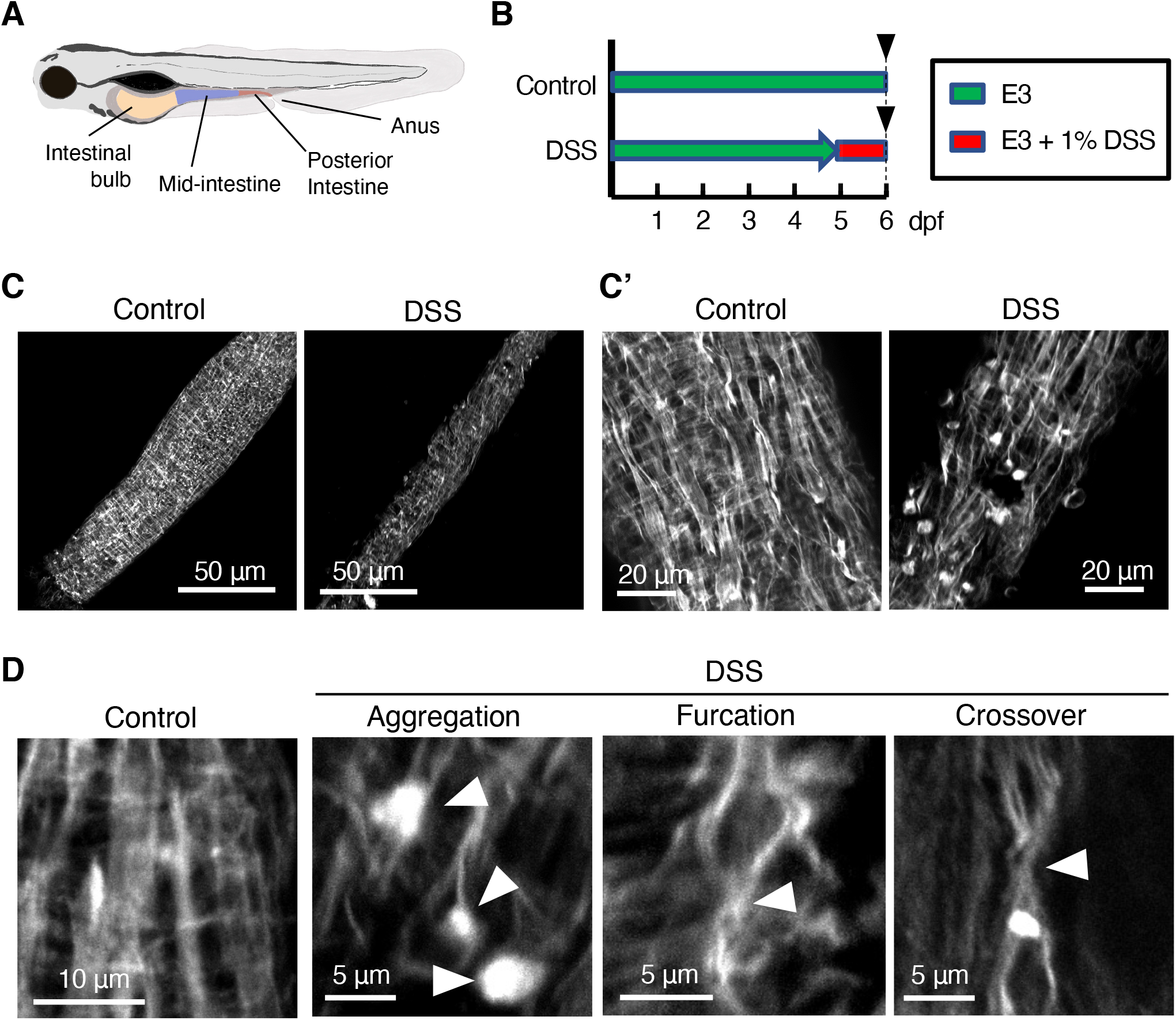
DSS administration damages the intestinal visceral musculature in zebrafish larvae. **A.** Schematic diagram of the larval zebrafish intestine showing three segments: intestinal bulb (yellow), mid-intestine (purple) and posterior intestine (red). Mid- and posterior intestine were observed for this study. **B.** The experimental schematic. Arrowhead indicates time of dissection **C.** The visceral musculature at 6 days of post-fertilization (dpf). The samples were stained by Phalloidin-594. Closer views of the visceral musculature are in C’. **D.** Representative visceral musculature disruption phenotypes.

## Discussion

Although the development of the visceral musculature has been studied in multiple model organisms (Aghajanian et al., 2016; Gabella and Clarke, 1983; Rudolf et al., 2014; Whitesell et al., 2014; Yang et al., 1999), the mechanisms by which the visceral musculature degenerates are poorly understood. Our study demonstrates that intrinsic and extrinsic factors can disrupt the visceral musculature, producing four prominent visceral musculature disruption phenotypes—sprout, discontinuity, furcation, and crossover (Figure 7A)—which can be used as quantitative readouts of disruption of the visceral musculature. Interestingly, the occurrence of sprout and discontinuity are significantly increased in all the treatment conditions compared to controls (Figure 7B). Thus, we speculate that sprout and discontinuity can be induced by mild stress to the visceral muscle, such as diet change and aging (Figure 7B). In contrast, the occurrence of furcation and crossover are not altered by *yki^3S/A^*-induced intestinal epithelial hyperplasia, diet change, or aging and only slightly induced even by *Ras^V12^* expression in ISCs and EBs or bleomycin feeding (Figure 7B). These data suggest that more severe perturbations are required for inducing the furcation and crossover phenotypes. Dissecting and staining the midguts to visualize the visceral musculature is quite straightforward, and these phenotypes are readily quantifiable. Therefore, our study provides insights into the deterioration of the visceral musculature and establishes a framework for investigating the molecular mechanisms underlying the visceral musculature degeneration in *Drosophila* as well as other animals.

**Figure 7:**
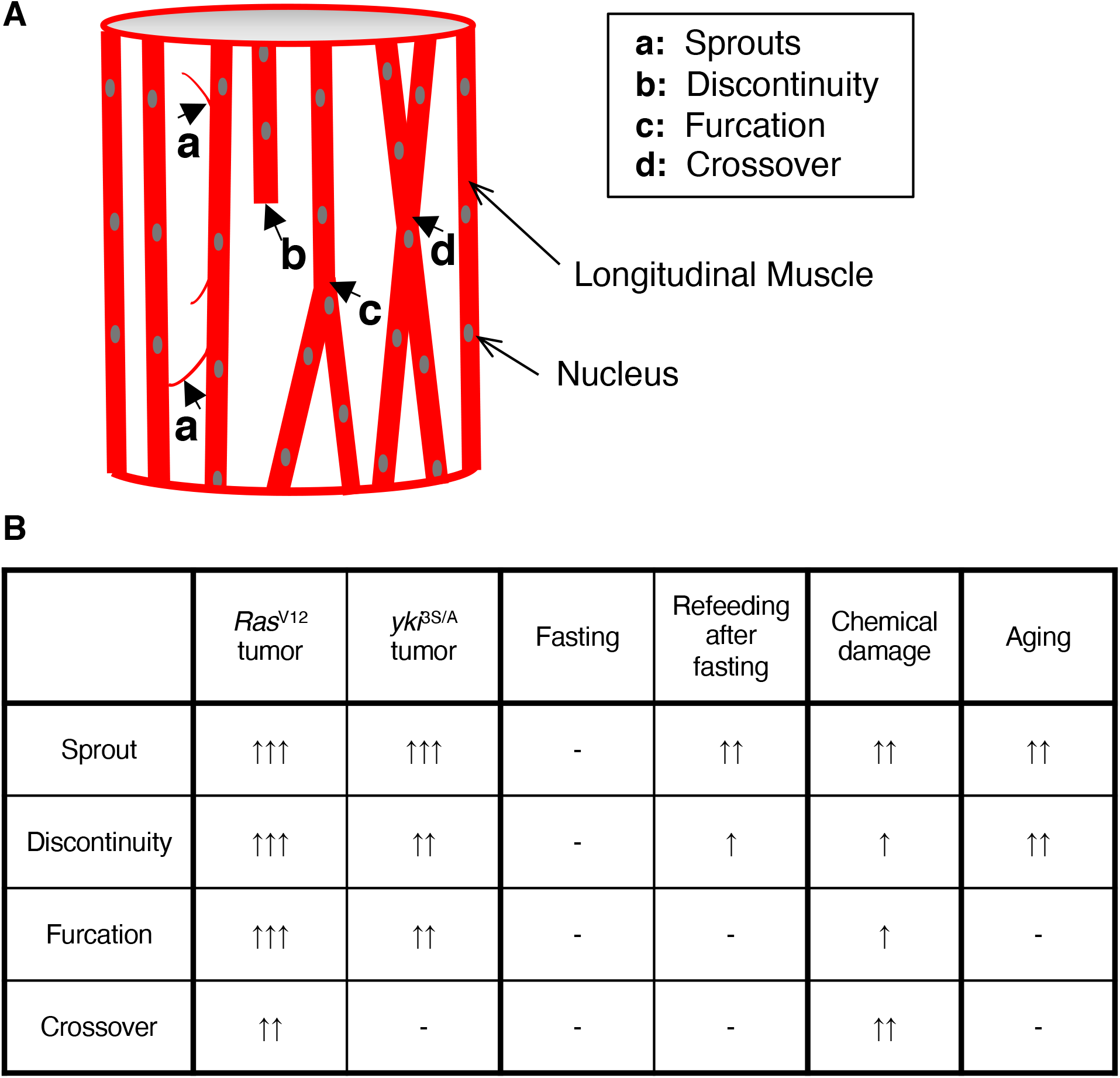
Summary of the visceral musculature disruption phenotypes in *Drosophila*. **A.** Diagram representing the visceral musculature disruption phenotypes. **B.** Summary of the occurrence of the visceral musculature disruption phenotypes under the tested experimental conditions. The data chosen for comparison is as follows: 4 day expression of *Ras*^V12^, 8 day expression of *yki*^3S/A^, 2 day fasting, 2 day re-feeding of rich diet for 2 days after fasting, 4 days on bleomycin, and 60 day old flies. (−) denotes < 50% increase, (↑) denotes 50-100% increase, (↑↑) denotes 100-400% increase, and (↑↑↑) denotes > 400% increase.

Our study also reveals an association between the visceral muscle and the neighboring intestinal epithelium. We show that oncogenic stress in the intestinal epithelium disrupts the visceral musculature (Figure 1 and 2). Although tension due to hyperplasia could be a factor that induces disruption of the visceral musculature, it is likely that additional factors associated with oncogenic transformation of epithelial cells are involved. For example, depletion of *Cortactin* specifically in *Ras^V12^*-expressing epithelial cells at least partially rescues the sprout and discontinuity phenotypes (Figure 1C), demonstrating that a process in intestinal epithelial cells can be a cause of the visceral musculature disruption. Bleomycin has been used in *Drosophila* to study how intestinal epithelium regenerates in response to tissue damage. Our findings indicate that bleomycin feeding also disrupts the visceral musculature. In addition to the role in peristalsis, the *Drosophila* visceral muscle has been shown to be a source for multiple signaling ligands, which play essential roles in the homeostasis and regeneration of the midgut epithelium (Biteau and Jasper, 2011; Guo et al., 2013; Jiang and Edgar, 2009; O’Brien et al., 2011; Scopelliti et al., 2014; Xu et al., 2011; Zhou et al., 2015). Therefore, it would be interesting to investigate whether disruption of the visceral musculature could influence the function and physiology of intestinal epithelium. Similarly, we show that DSS feeding also disrupts the visceral musculature in zebrafish larvae. Given that DSS administration is used to model IBD in zebrafish and other animal models (Chuang et al., 2019; Kiesler et al., 2015; Oehlers et al., 2013; Oehlers et al., 2017), our findings raise the possibility that disruption of the visceral muscle structure could be an aspect of IBD. It has been shown that gastrointestinal motility disorders are frequently associated with IBD (Bassotti et al., 2014; Hashmi et al., 2021). Therefore, it would be interesting to investigate whether the visceral musculature is affected in IBD animal models and patients.

*Drosophila* have been extensively used for aging research because of several advantages, including short lifespan, low cost, and powerful genetics. Multiple aging-associated phenotypes available in *Drosophila* allow the monitoring of organismal, tissue-level, and molecular aging processes (He and Jasper, 2014; Piper and Partridge, 2018). Although gradual deterioration of tissue structure is a feature of aging, directly assessing aging-related deterioration of a tissue structure and pattern is not straightforward in *Drosophila*. Our study demonstrates that the visceral musculature degenerates over time and also as the result of other intrinsic and extrinsic stresses, which can be quantified by the visceral musculature disruption phenotypes. Visceral muscle stem or progenitor cells have not been identified in adult *Drosophila*. Thus, the visceral muscle’s regeneration and repair capacity is expected to be limited. Our study raises a possibility of using the visceral musculature disruption phenotypes as markers for assessing wear and tear of the tissue structure during aging. Studies elucidating the inverse correlation between longevity and the occurrence of the visceral musculature disruption phenotypes will help to further test this possibility.

## Acknowledgements

This work was in part supported by R35GM128752 to Y.V.K. from the National Institutes of Health.

## Materials and Methods

### Fly genetics and husbandry

Flies were maintained with standard cornmeal-agar medium at room temperature. 4-day-old non-virgin female flies were collected and used for experiments. For *Ras*^V12^ and *yki*^3S/A^ overexpression assay, flies were reared at 18°C, and the collected flies were placed at 29°C for indicated time period to induce transgenes (Kwon et al., 2015; Lee et al., 2020). During 29°C incubation, flies were transferred to fresh food vials every 2 days.

For the starvation, chemical damage, and aging experiments, we used *w^1118^* (#3605) obtained from Bloomington Drosophila Stock Center (BDSC). To induce ISCs and EB cell dissemination, we used *esg-GAL4*, *tub-GAL80^ts^*, *UAS-GFP* (referred as *esg^ts^*) crossed with one of the following: *UAS-yki^3S/A^* (BDSC #28817), *UAS-Ras^V12^* (laboratory stock), and *UAS-Cortactin RNAi* (HMS00658; BDSC #32871).

### Starvation assay

Starvation experiment was adapted from Lucchetta et al (Lucchetta and Ohlstein, 2017). The flies were maintained on a rich diet (standard molasses food supplemented with live yeast mixed with 5% sucrose) for 2 days. The flies were then divided into 3 cohort groups (10 flies for each). Group 1 was fed on the rich diet for an additional 2 days. Groups 2 and 3 were maintained in vials containing a piece of 2.5 × 3.75 cm^2^ chromatography paper soaked with 0.5 ml of deionized water for 2 days for starvation. Group 3 was then fed on the fresh rich diet for 2 days.

### Bleomycin feeding assay

Bleomycin administration procedure was adapted from Amcheslavsky et al. (Amcheslavsky et al., 2009). Flies were divided into 2 cohort groups (10 flies each). Group 1 was maintained in a vial containing a piece of 2.5 × 3.75 cm^2^ chromatography paper soaked in 0.5 ml of 5% sucrose for 4 days and served as controls. Group 2 was maintained in a vial with a piece of 2.5 × 3.75 cm^2^ chromatography paper soaked in 0.5ml solution of 125 μg/ml bleomycin (Sigma, B1141000) in 5% sucrose for 4 days.

### Aging experiment

15 flies for each aging group were maintained at room temperature on a standard molasses food for the indicated duration of the experiment. Flies were transferred to fresh food every 5 days.

### Immunohistochemistry

Female flies were dissected in Phosphate Buffered Saline (PBS). Dissected guts were fixed in 4% paraformaldehyde (Electron Microscopy Services, RT15710) diluted in PBS for 20 minutes at room temperature. The fixed guts were then washed two times with PBST (PBS containing 0.2% Triton-X). For permeabilization and blocking, we incubated the tissue samples in a blocking buffer (PBST supplemented with 5% normal goat serum) for 1 hour at room temperature. The tissue samples were then incubated with Phalloidin conjugated to Alexa Fluor 594 (1:1000) and DAPI (1:2000) for 1 hour at room temperature or overnight at 4°C. The guts were washed with PBST and mounted with Vectashield mounting medium (Vector Laboratories, H-1000).

### Confocal microscopy

Midguts were imaged at 40x/1.25 oil objective lens using Leica SP8 laser scanning confocal microscope. Images were processed using FIJI ImageJ software, with Z-projections averaging 2-3 μm thickness allowing most longitudinal musculature to show in a 388μm x 388μm confocal microscope field. The posterior midgut of *Drosophila* was chosen for imaging and quantification.

### Quantification of VM Damage

The VM disruption phenotypes were counted from one leaflet of the R5 region of the posterior midgut captured in 388 μm × 388 μm confocal microscope field. The four visceral muscle disruption phenotypes were defined and then quantified as in number of events per posterior midgut image within the field. For sprouts, each continuous fibril that emerges from the main longitudinal muscle was considered as a single sprout. For discontinuity, every non-continuous stub observed within the field of view was considered as an event. For furcation, every point that resulted in the change in the number of continuous main longitudinal muscles due to merging/diverging was considered as a event. For crossover, every point where two main longitudinal muscles overlapped without merging was considered as a event.

### Zebrafish husbandry and maintenance

Adult zebrafish were maintained at 28.5 °C under a 16h:8h light:dark cycle. Fish stocks of WT(ABb), a derivative of wild-type AB^wp^, were used to obtain embryos. Embryos were collected in E3 medium (5.0mM NaCl, 0.17mM KCl, 0.33mM CaCl_2_, 0.33mM MgCl_2_·6H_2_O, adjusted to pH7.2-7.4) in petri dishes by in vitro fertilization as described in Westerfield et al. (Westerfield, 2007) with modifications. Unfertilized and dead embryos were removed 5 hours post-fertilization (hdf) and 1 day post-fertilization (dpf). Fertilized embryos were kept in E3 at 28.5°C until 5 dpf and 6 dpf for use in experiments. All animal work in this study was conducted with approval of the University of California Irvine Institutional Animal Care and Use Committee (Protocol #AUP-19-043) in accordance with the institutional and federal guidelines for the ethical use of animals.

### Zebrafish Dextran Sulfate Sodium (DSS) overnight treatment

We adapted and modified the DSS overnight treatment protocol described previously (Chuang et al., 2019; Oehlers et al., 2013). Embryos were raised in 10ml E3 until 5 days post-fertilization (dpf), at which time 15 larvae were transferred to freshly prepared 1% (w/v) DSS (SIGMA, D8906-10G, MW>500,000) in E3 for a single overnight treatment. Larvae were not fed on the DSS treatment day.

### Zebrafish Dissection, Fixation, Staining and Imaging

Larval dissection protocol to obtain intestines was adapted and modified from San et al. (San et al., 2018). 15 control larvae were raised in 10ml E3 and dissected on 6 dpf. 15 experimental larvae were raised in 10ml E3 and were transferred to freshly prepared 1% DSS on 5 dpf and dissected on 6 dpf. Dissection was performed under a Zeiss SteREO Discovery V20 Microscope. 0.4% (w/v) tricaine stock (pH 7-7.5) was diluted to a working concentration of 0.1% (v/v) in E3. Up to 3 larvae were placed into anesthetic and each larva was transferred to a petri dish lid under the microscope for dissection. A precision watchmaker’s forceps was used to stabilize the fish body whilst a micro-dissecting scissor was used to remove the fish head to expose the intestine. Another precision watchmaker’s forceps was then used to pinch the swim bladder to remove the intestine in the anterior direction. Extra tissues were removed gently with both forceps and intestines were placed in 4% paraformaldehyde for a 2 hour fixation as described in Wallace et al. (Wallace et al., 2005) followed by three 5-minute washing steps in 1x PBST. Fixed intestines were placed in phalloidin 594 (1:400 in blocking buffer) for 1h before proceeding to another three 5-minute washing steps in 1x PBST. Intestines were then mounted and imaged with 40X water-immersion objective on a Leica SP8 confocal microscope.

### Statistics and reproducibility

All the images presented and used for quantification are from the posterior R5 region of adult female fly midguts. All experiments were independently repeated at least three times. Statistical analyses were performed using Microsoft Excel. All *P* values were determined by two-tailed Student’s t-test with unequal variances. Level of significance are depicted by asterisks in the figures: \**P*< 0.05; \*\**P*< 0.01; \*\*\**P*< 0.001. Sample sizes were chosen empirically based on the observed effects and indicated in the figure legends.

