## Supplemental Figures for "Wear and Tear of the Intestinal Visceral Musculature by Intrinsic and Extrinsic Factors"

### Supplemental Figure 1

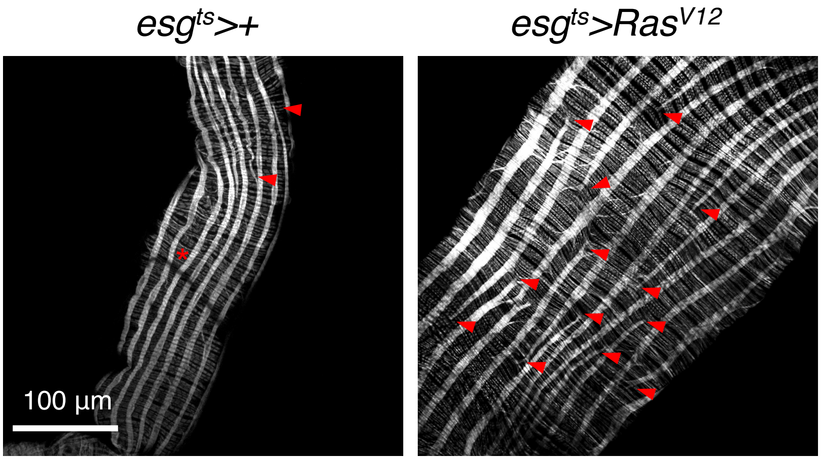


Surface views of the posterior midgut. Visceral musculature (gray) is visualized with Phalloidin staining. Arrowheads show discontinued longitudinal muscles. The region void of Phalloidin signals (asterisks) is where the trachea is present. Muscles behind the trachea are not captured because they are at different focal planes. Scale bar, 100 $\mu$m.

#### Supplemental Figure 2

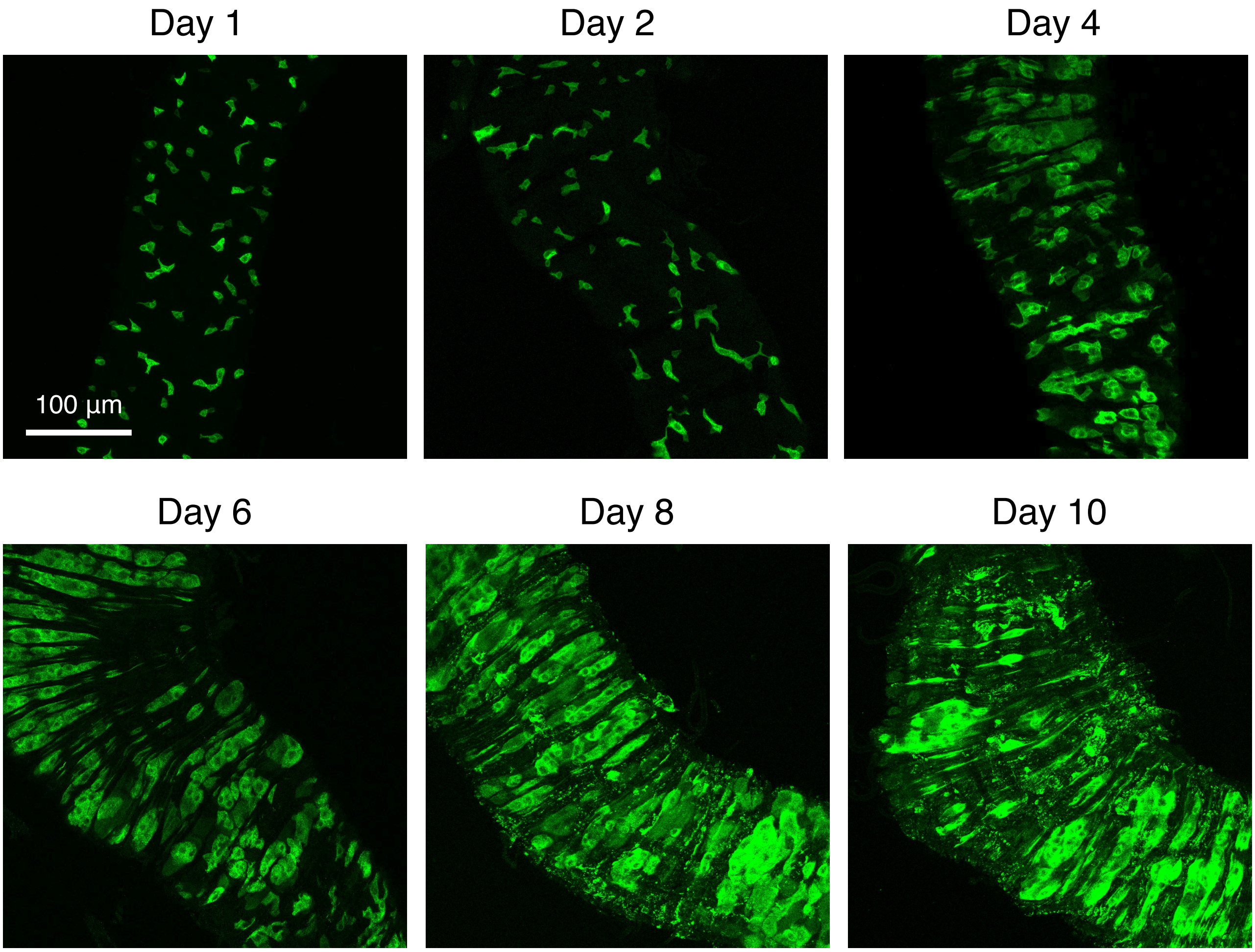


Images of the *$esg^{ts}>yki^{3S/A}$* posterior midgut. Transgenes were induced with *$esg^{ts}$* by incubating at 29°C for indicated durations. The cells manipulated by *$esg^{ts}$* are marked and stained with GFP (green). Scale bar, 100 $\mu$m.
